# Morphological innovation and lineage-specific history drive disparification in the aggregated pollen of mimosoid plants

**DOI:** 10.1101/2025.04.16.649038

**Authors:** Rafael F. Barduzzi, Stefany Liau-Kang, Ana Flávia Trabuco Duarte, Francisco de Assis Ribeiro dos Santos, Leonardo M. Borges

**Affiliations:** Departamento de Botânica, Universidade Federal de São Carlos, São Carlos, Brazil; Faculdade de Filosofia, Ciências e Letras de Ribeirão Preto, Universidade de São Paulo, Ribeirão Preto, Brazil; Universidade Estadual de Campinas, Campinas, Brazil; Departamento de Ciências Biológicas, Universidade Estadual de Feira de Santana, Feira de Santana, Brazil

**Keywords:** Caesalpinioideae, disparity, ecological groups, Fabaceae, Leguminosae, morphological diversity, Mimosoid clade, palynology, phylomorphospace, Stryphnodendron clade

## Abstract

**Background and Aims:** The study of morphological diversity (i.e., disparity) offers unique opportunities to understand evolutionary patterns and processes. Plant disparity studies reveal that morphological disparification can be related to factors such as secondary woodiness or to pollination niche, for example. Similarly, some pollen traits are known to be shaped by environmental pressures, but this influence has only been evaluated in monads, never in multi-grained dispersal units. In this study, we investigated the disparity of aggregated dispersal units in two lineages of Neotropical mimosoid legumes. The Mimosa and Stryphnodendron clades are independent lineages that share similarities in pollen morphology and biome shifts. In this context, we asked: What are the patterns of pollen disparity in these lineages, and are these patterns similar between lineages occurring in the same biomes?

**Methods:** To answer these questions, we compiled data from the literature on pollen morphology and biomes of occurrence for a phylogenetically representative set of taxa in the Mimosa and Stryphnodendron clades. With these data, we calculated morphospaces and disparity metrics, and tested whether the pollen morphology of distinct lineages occurring in the same biome differs significantly.

**Key Results:** Our results show that Mimosa and Stryphnodendron clades exhibit distinct patterns of pollen disparity, as do independent lineages occurring in the same biomes. Additionally, we observed that certain biomes support greater or lesser levels of morphological disparity.

**Conclusions:** We conclude that (1) the Mimosa clade has greater disparity, possibly due to evolution of novel pollen morphologies in the genus *Mimosa*, (2) there is a maintenance of similarities in the pollen of the Stryphnodendron clade, *Adenopodia* and *Piptadenia*, and (3) the evolution of pollen grains in these groups appears to be primarily shaped by phylogeny and developmental constraints, with environmental pressures playing a comparatively smaller role.

## INTRODUCTION

The study of morphological diversity (i.e., disparity) offers unique opportunities to understand how the interaction between morphology and environment shapes plant evolution. For example, the shift from herbaceous to woody habit promoted disparification of different plant lineages after colonization of insular habitats (Nürk *et al*., 2019). Similarly, in Brassicaceae, variation in floral disparity follows changes in pollination niche (Gómez *et al*., 2016). However, most studies on plant disparity focus solely on macromorphology, or use traits such as plant height as proxies for overall morphological diversity (e.g., Chartier et al., 2014; Xue *et al*., 2015; Gómez *et al*., 2016, 2022; Nürk *et al*., 2019). The disparity of microcharacters such as those in pollen grains is rarely investigated (e.g., Mander, 2016; Kriebel *et al*., 2017; Jardine *et al*., 2022), leaving open questions about the influence of environmental factors over the evolution of traits with prominent reproductive and ecological importance.

Pollen grains are naturally able to change their shape, particularly by folding in response to dehydration (Katifori *et al*., 2010). Thus, pollen evolution should be sensitive to environmental pressures. Indeed, studies focusing on specific traits demonstrate that environmental conditions shape pollen structure, composition, and physiology (Pacini and Franchi, 2020). For instance, variations in wall ornamentation may be linked to different pollination mechanisms (Osborn *et al*., 1991; Banks and Rudall, 2016). Moreover, a reduction or complete loss of the exine layer is commonly observed in aquatic angiosperms (Osborn *et al*., 2001), and the presence of two types of furrows in Acanthaceae and Boraginaceae may be an adaptation to dry environments (Pacini and Franchi, 2020). Similarly, one of the few studies on pollen disparity highlights the influence of variations in moisture and temperature across the latitudinal gradient over the morphology of Myrtales pollen (Kriebel *et al*., 2017).

However, environmental pressures that apply to single pollen grains (monads) may not equally affect pollen organized in multi-grained dispersal units (dyads, tetrads or polyads). As such dispersal units are commonly dispersed dehydrated and do not break apart after release from the anther, they probably have adaptations to environmentally-driven dehydration that could cause the grains to fold and separate (Franchi *et al*., 2002). In polyads, for example, variations in aperture morphology and continuity of tectum have enabled individual monads to respond to dehydration as a single unit (Banks *et al*., 2010; Elazab, 2016). In addition to surface exposure, other differences in the development and morphology of aggregated units may also change how they interact with the environment (Banks *et al*., 2010). Hence, similar environmental pressures may have led to different morphological adaptations between monads and aggregated units.

Aggregated dispersal units are particularly common in mimosoid legumes (former subfamily Mimosoideae, now tribe Mimoseae of the Caesalpinioideae; Bruneau *et al*., 2024). The shape and size of pollen in the tribe are known to vary widely, showcasing both the largest and smallest aggregated units of all angiosperms (Banks *et al*., 2010). Two Mimoseae lineages, the Mimosa and Stryphnodendron clades (Borges, Simon, Morales, *et al*., 2024; Borges, Simon, Ribeiro, *et al*., 2024), include species with tetrads and polyads (Fig. 1), features that were once used as evidence for including them in the same informal group (the Piptadenia group; Lewis and Elias, 1981). Together, these clades include ten genera (*Adenopodia* C. Presl, *Gwilymia* A.G. Lima, Paula-Souza & Scalon, *Marlimorimia* L.P. Queiroz, L.M. Borges, Marc.F. Simon & P.G. Ribeiro, *Microlobius* C. Presl, *Mimosa* L., *Naiadendron* A.G. Lima, Paula-Souza & Scalon, *Parapiptadenia* Brenan, *Piptadenia* Benth., *Pityrocarpa* (Benth.) Britton & Rose, *Stryphnodendron* Mart.) that exhibit relatively wide pollen diversity (Guinet and Caccavari, 1992; Caccavari, 2002; Buril *et al*., 2010; Santos-Silva *et al*., 2013; Ribeiro *et al*., 2018) and share similar evolutionary histories. These histories include parallel biome shifts between seasonally dry tropical forest (hereafter “SDTF“; e.g., Caatinga), tropical rainforest (e.g., Atlantic Forest and Amazon Forest), and savanna (e.g., Cerrado) (Simon *et al*., 2016; Ribeiro *et al*., 2018; Borges *et al*., 2022; Lima *et al*., 2022; Ringelberg *et al*., 2023). Sharing similar lineage biome shifts and variation in pollen morphology in these two independent lineages (Fig. 1) makes the Mimosa and Stryphnodendron clades interesting models to investigate if and how habitat shapes pollen disparity across time.

**Fig. 1.**
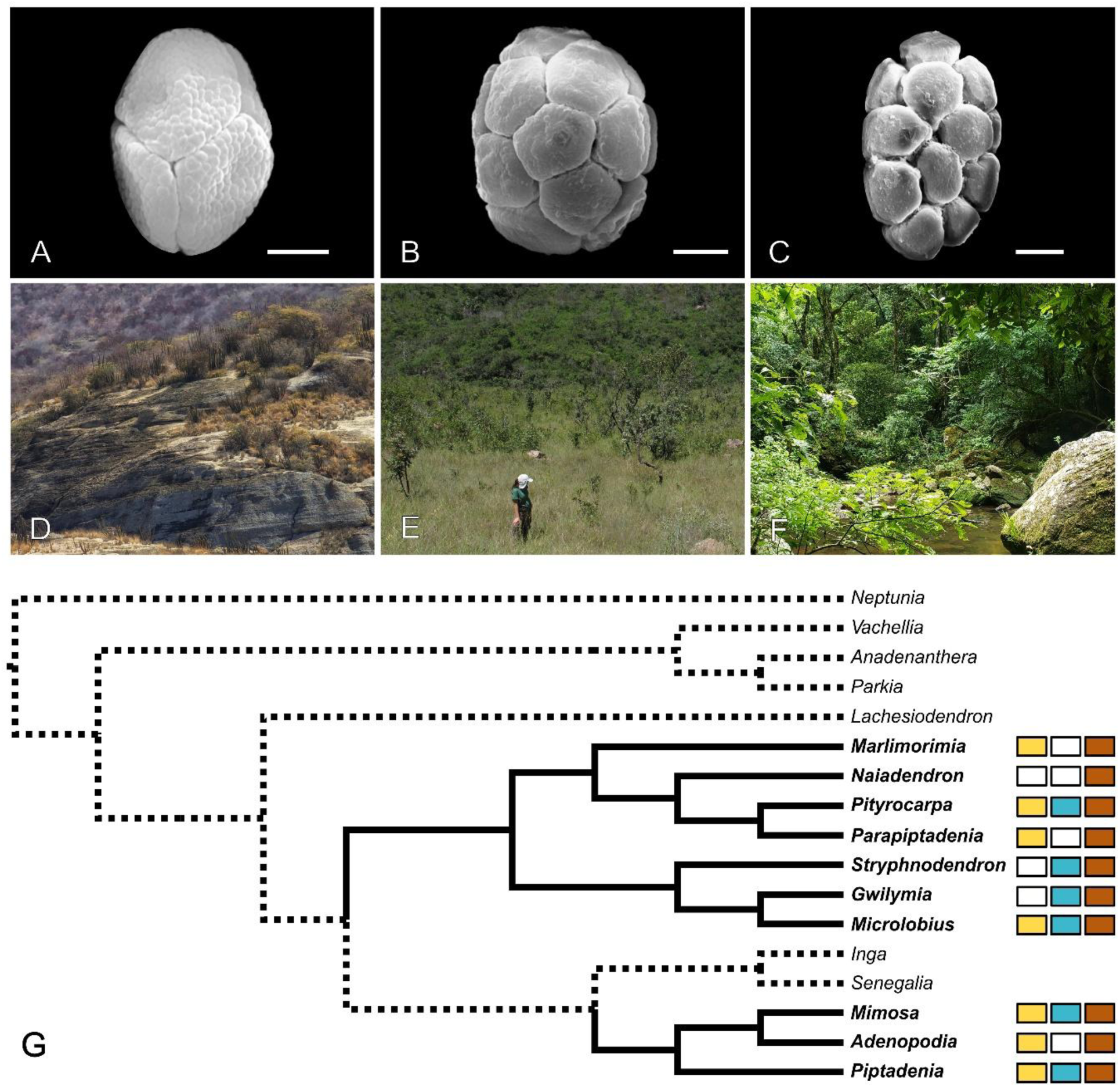
(A–C) Examples of pollen morphologies in the Mimosa and Stryphnodendron clades. (A) Tetrad of *Mimosa oedoclada* Barneby. (B) Polyad of *Parapiptadenia rigida* (Benth.) Brenan. (C) Polyad of *Stryphnodendron adstringens* (Mart.) Coville. (D–F) Biomes where species of Mimosa and Stryphnodendron clades occur. (D) Succulent. (E) Savanna. (F) Rainforest. (G) Phylogenetic relationships among some mimosoid legumes (modified from Hughes *et al*., 2022). Lineages in black belong to the Mimosa and Stryphnodendron clades. Boxes indicate biomes of occurrence: Yellow: Seasonally dry tropical forest, Blue: Savanna, Brown: Rainforest. Images: (A-C) S. Liau-Kang. (D) O. Nogueira. (E) L. M. Borges. (F) K. Nagata. Scale bars: 5μm.

Here, we aimed to identify how pollen grain disparity varies with the biome of occurrence. Therefore, we asked: (1) What are the patterns of pollen disparity in different lineages of the Mimosa and Stryphnodendron clades?, (2) Does pollen disparity vary across different biomes?, and (3) Do independent lineages inhabiting the same biomes exhibit similar patterns of pollen disparity? As environmental pressures may constrain pollen diversification, we expected that lineages occurring in similar biomes would have similar disparity patterns.

## MATERIALS AND METHODS

### Data sampling and treatment

We compiled pollen morphological data from the literature (Barth and Yoneshigue, 1966; Barth, 1973, 1974; Kenrick and Knox, 1982; Guinet and Caccavari, 1992; Caccavari, 2002; Flores-Cruz *et al*., 2006; Lima *et al*., 2008; Bauermann *et al*., 2009; Braga *et al*., 2012; Santos-Silva *et al*., 2013; Ribeiro *et al*., 2018; Cruz *et al*., 2018; Medina-Acosta *et al*., 2019; Barduzzi *et al*., 2024; Liau-Kang *et al*., 2024). In cases of polymorphic characters described with more frequent and less frequent states, we included only the most frequent trait variations. Terminology used in different literature sources was standardized according to Punt *et al*. (2007) and Halbritter *et al*. (2018) to ensure consistency.

Our sample included 193 taxa representing species, subspecies and varieties from all genera in the Mimosa and Stryphnodendron clades, plus the three closely related genera *Inga* Mill., *Lachesiodendron* P.G. Ribeiro, L.P. Queiroz & Luckow, and *Senegalia* Raf. Although the initial character sample included 11 characters, we removed four due to their low completeness (fewer than 50%) to avoid biases due to high proportions of missing data (Lehmann *et al*., 2019). The final phenotypic sample included seven characters, four categorical and three continuous (Table 1).

**Table 1.** Pollen characters evaluated. Character states are provided for categorical and discrete traits, while value ranges are reported for continuous traits.

| Character | States |
| --- | --- |
| Dispersal unit - number of grains | (0) one; (1) two; (2) four; (3) eight; (4) twelve; (5) sixteen; (6) thirty two |
| Dispersal unit - outline | (0) circular; (1) elliptical; (2) oval |
| Pollen grains arrangement | (0) decussate; (1) irregular arrangement polyad; (2) regular arrangement polyad; (3) rhomboidal; (4) tetragonal; (5) tetrahedral |
| Pollen ornamentation | (0) areolate; (1) fossulate; (2) microverrucate; (3) rugulate; (4) psilate; (5) reticulate; (6) verrucate; (7) verrucate-scabrate; (8) scabrate |
| Dispersal unit shorter diameter | 8.3 - 50µm |
| Dispersal unit longer diameter | 8.6 - 100µm |
| Exine thickness | 0.15 - 2.5µm |

We compiled information on the vegetation type and/or domain of occurrence of each taxon using data available in the Flora do Brasil website (Brazil Flora Group, 2022) and in taxonomic literature (Barneby, 1991; Simon *et al*., 2009; Ribeiro *et al*., 2018; Borges *et al*., 2022; Lima *et al*., 2022). Data from the Brazilian Flora website were downloaded using the “flora” package (Carvalho, 2020) in R (R Core Team, 2014). Based on this dataset, we determined the occurrence of each taxon in six biomes: desert, flooded grassland, savanna, SDTF, temperate and tropical rainforest.

### Phylogenetic hypothesis

A phylogenetic tree is necessary for the disparity analyses used here (see below). However, current phylogenetic analyses that include both the Mimosa and Stryphnodendron clades have sparse taxon sampling (Koenen *et al*., 2020; Ringelberg *et al*., 2023). To overcome this problem, we synthesized the topology of the most taxonomic comprehensive analyses for these two clades to date (Vasconcelos *et al*., 2020; Borges *et al*., 2022; Lima *et al*., 2022). We then pruned the tree to match the morphological dataset and set branch lengths to one to avoid biases related to variation in branch length estimation in the original analyses. Tree editing was performed using the package “ape” (Paradis and Schliep, 2019).

### Pollen morphospaces

Variation in pollen morphology among members of the Mimosa and Stryphnodendron clades was evaluated using a morphospace. To achieve this, we first estimated the optimal number of intervals describing the distribution of each continuous character using the Freedman-Diaconis (F-D) rule for histograms (Freedman and Diaconis, 1981). Then, we discretized the characters with the function “discretizeDF” from the “arules” package (Hahsler *et al*., 2005). Discretization was necessary because available packages for morphospace estimation do not allow joint use of categorical and continuous variables present in our dataset. Later, with the “Claddis” package (Lloyd, 2016), we performed a Principal Coordinate Analysis (PCoA) based on a distance matrix produced with the Maximum Observable Distance (MORD; Lloyd, 2016). All characters were treated as unordered and as having equal unit weight to avoid biases related to the evolutionary direction or taxonomic importance of characters (Congreve and Lamsdell, 2016; Bansal *et al*., 2021). Polymorphisms were resolved by selecting the character state generating the smallest distance. To avoid negative eigenvalues, the PCoA was carried out using the Cailliez correction. Finally, we plotted the morphospace using the first two PCoA axes, superimposing the phylogeny to highlight lineages (i.e., phylomorphospace) or to delimit convex hulls based on biomes of occurrence to highlight ecological groups.

### Ecological lineages delimitation

Apart from investigating the disparity of clades and ecological groups, we delimited “ecological lineages” to compare disparity between independent lineages occurring in the same biomes. Using the complete phylogeny for the Mimosa and Stryphnodendron clades, we inferred ancestral habitats with the stochastic mapping function (“make.simmap”) of the “ape” package. Then, we searched for clades of three or more species with node likelihood above 50% for one particular biome to delimit ecological lineages. Finally, for delimited lineages with available pollen data, we applied the analysis described below.

### Disparity metrics, overlap, and statistical analyses

We used the “Claddis” and “dispRity” packages (Lloyd, 2016; Guillerme, 2018) to calculate three disparity metrics: mean pairwise distance (MPD), sum of variances (SV) and sum of ranges (SR). MPD characterizes density of morphospace occupancy and was calculated from the distance matrix. SV and SR describe the extent of the morphospace occupancy based on all PCoA axes and were calculated both before and after 1000 bootstrap replications for each group (Guillerme *et al*., 2020). We calculated these metrics for both the Mimosa and Stryphnodendron clades, as well as for all genera and ecological groups and lineages. Metrics could not be calculated for genera lacking the minimum number of observations needed for analyses (two for the density metrics and three for the extent metrics). To evaluate the probability of overlap between morphological disparity calculated for different groups, we applied the Bhattacharyya coefficient (Bhattacharyya, 1943) to bootstrapped SV and SR values. We tested for statistically significant differences between SV and SR medians for all groups using the Wilcoxon rank-sum test (Wilcoxon, 1992).

## RESULTS

### Morphospace occupancy

The morphospace reveals an unequal disparity between the Mimosa and Stryphnodendron clades (Fig. 2). Taxa belonging to the genus *Mimosa* occupy a distinct part of the morphospace, while its close genera, *Adenopodia* and *Piptadenia*, are dispersed in a cluster together with members of the Stryphnodendron clade. Nonetheless, the distribution of each genus within this latter cluster is not homogeneous, as evidenced by the different areas occupied by *Gwilymia* and *Parapiptadenia* taxa, for example.

**Fig. 2.**
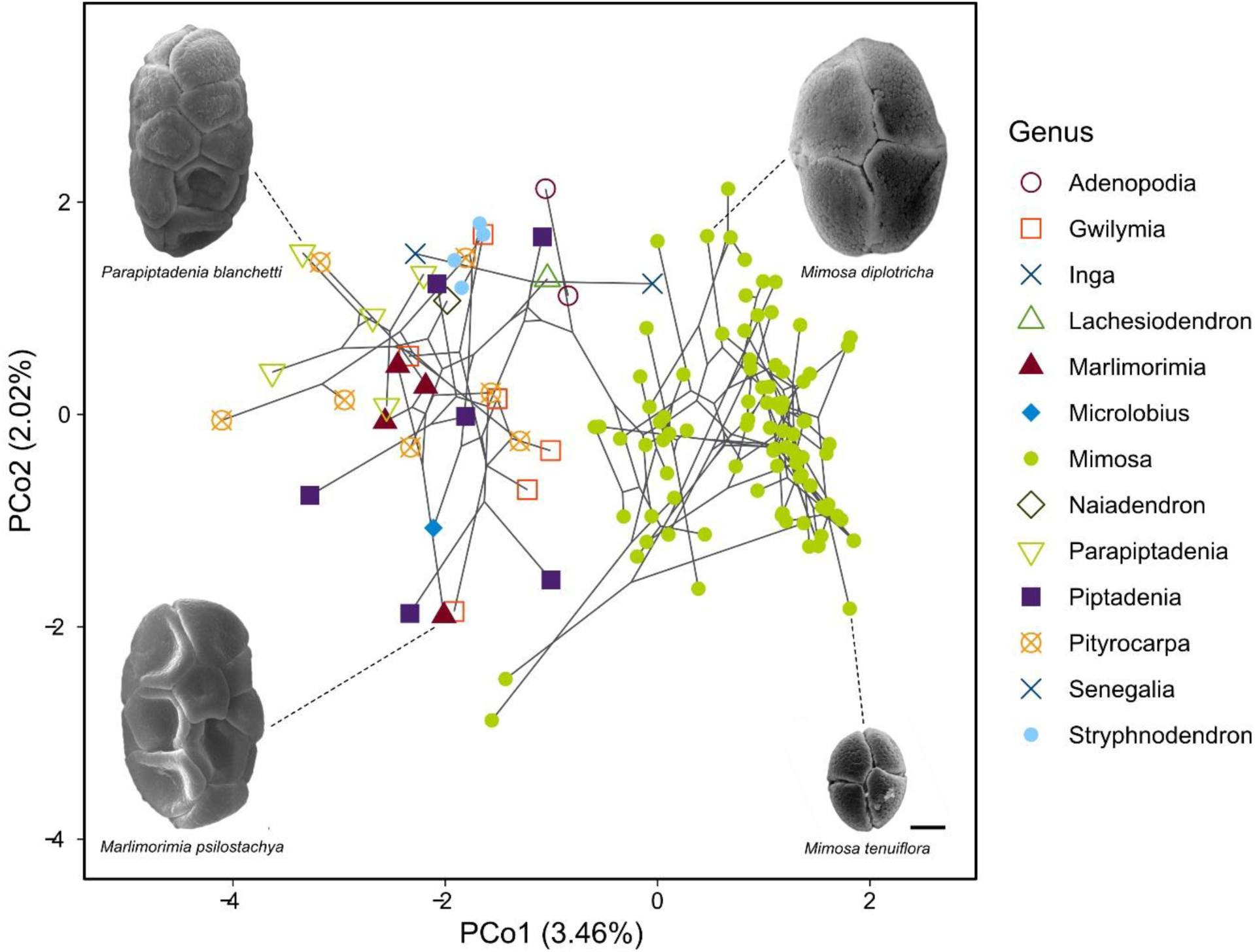
Phylomorphospace of Mimosa and Stryphnodendron clades, including the outgroups *Inga*, *Lachesiodendron*, and *Senegalia*. Values in parentheses indicate the percentage of variance explained by each axis. The folded grains in *Parapiptadenia blanchetti* and *Marlimorimia psilostachya* are a result from specimens dehydration and should not be accounted as morphological variation. Scale bar: 5μm.

### Morphological disparity of clades

Given the strong correlation between MPD and SV observable values (Spearman’s *rho* = 1, p < 1×10^−20^) and the fact that SR values merely emphasize the distinctions already evident in SV results (Supplementary Fig. S1; Table S1), we focus on the SV results here. Disparity metrics indicate that the Mimosa clade (represented by genus *Mimosa* and *Piptadenia* in our analysis) exhibits greater morphological disparity, occupying a larger extent of the morphospace (SV values; Fig. 3A; Supplementary Table S2) with a less dense distribution compared to the Stryphnodendron clade (MPD values; Supplementary Table S3). Within the Stryphnodendron clade, *Gwilymia* and *Pityrocarpa* seem to explore the morphospace further than *Marlimorimia, Parapiptadenia* and *Stryphnodendron* (Fig. 3B).

**Fig. 3.**
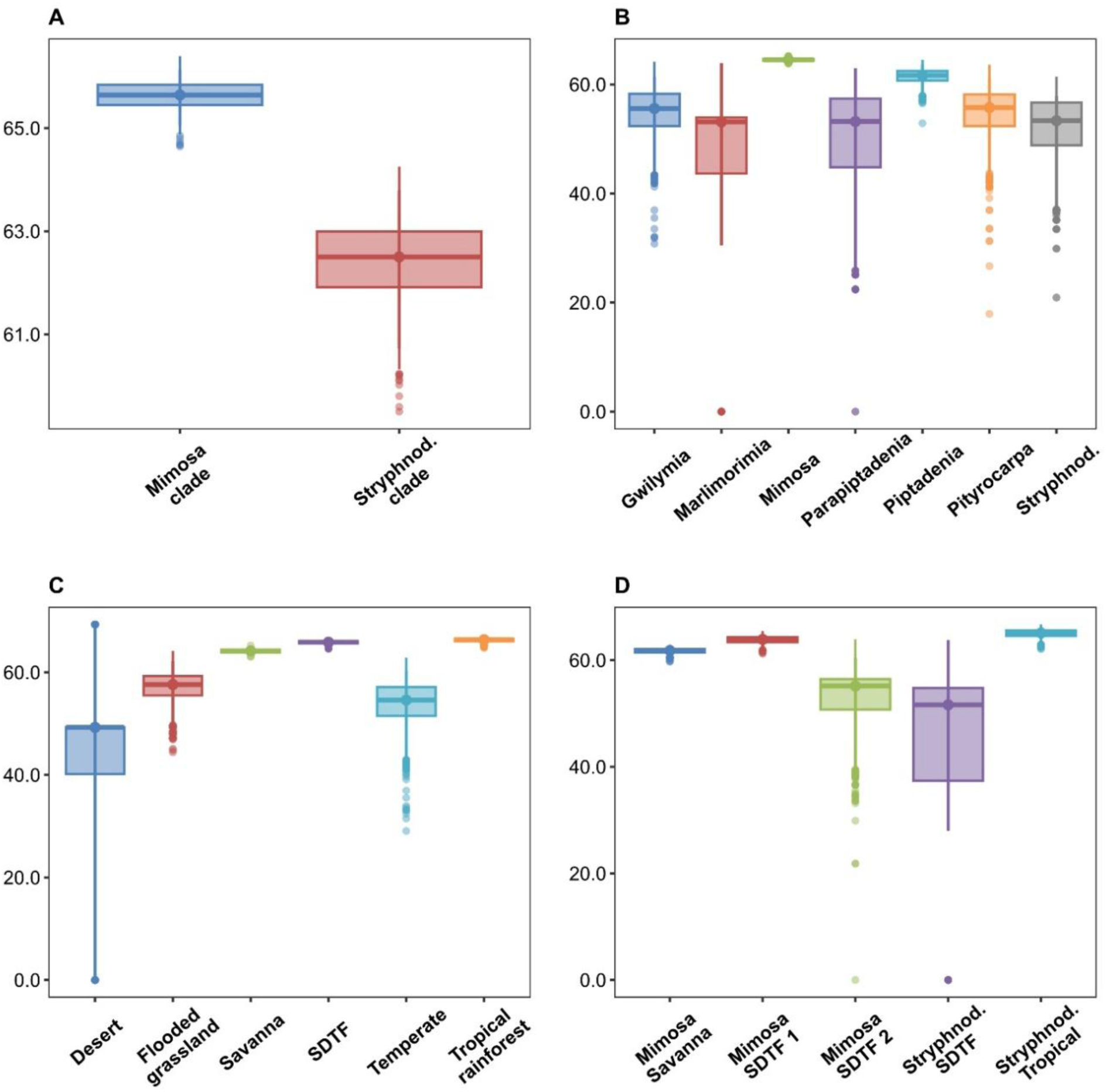
Morphological disparity (sum of variances) for groups. Comparisons between Mimosa and Stryphnodendron clades (A), genera (B), ecological groups (C), and ecological lineages (D). Boxplots show the distribution of values from 1000 bootstrap replicates.

The Bhattacharyya coefficient indicated that most Stryphnodendron clade genera have high probabilities of overlap (>50%) in their values for morphospace extent, with comparisons involving *Marlimorimia* being the only exceptions (Fig. 4A; Supplementary Table S4). In line with this, the Wilcoxon rank-sum test p-value indicates that *Parapiptadenia* and *Stryphnodendron* share significantly similar morphospace extents, a condition also seen for *Gwilymia* and *Pityrocarpa*. On the other hand, within the Mimosa clade, *Piptadenia* has lower chances of overlapping (<50%) in extent values with all Stryphnodendron clade genera, while *Mimosa* shows low overlap with any other group, occupying a dissimilar, greater morphospace extent (Fig. 4A).

**Fig. 4.**
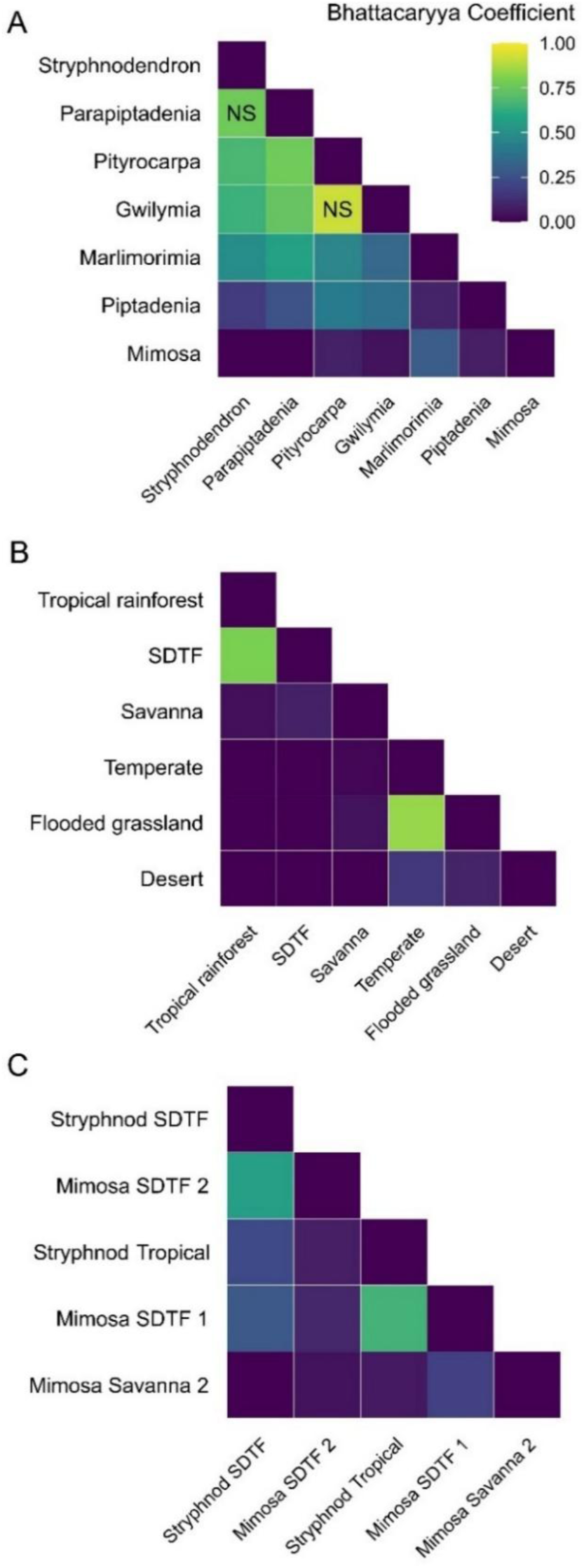
Dissimilarity matrix of Bhattacharyya values for sum of variances. ns: non-significant difference in disparity indicated by the Wilcoxon rank-sum test.

### Biomes space coverage and similarities between ecological lineages

The morphospace with convex hulls for ecological groups shows a high overlap between occupied morphospace areas, even though the limits of morphologies occurring in each biome are different (Fig. 5). Tropical rainforest, savanna, and SDTF are the biomes with the widest convex hulls, while flooded grassland, temperate and desert biomes appear to include a smaller diversity of pollen forms.

**Fig. 5.**
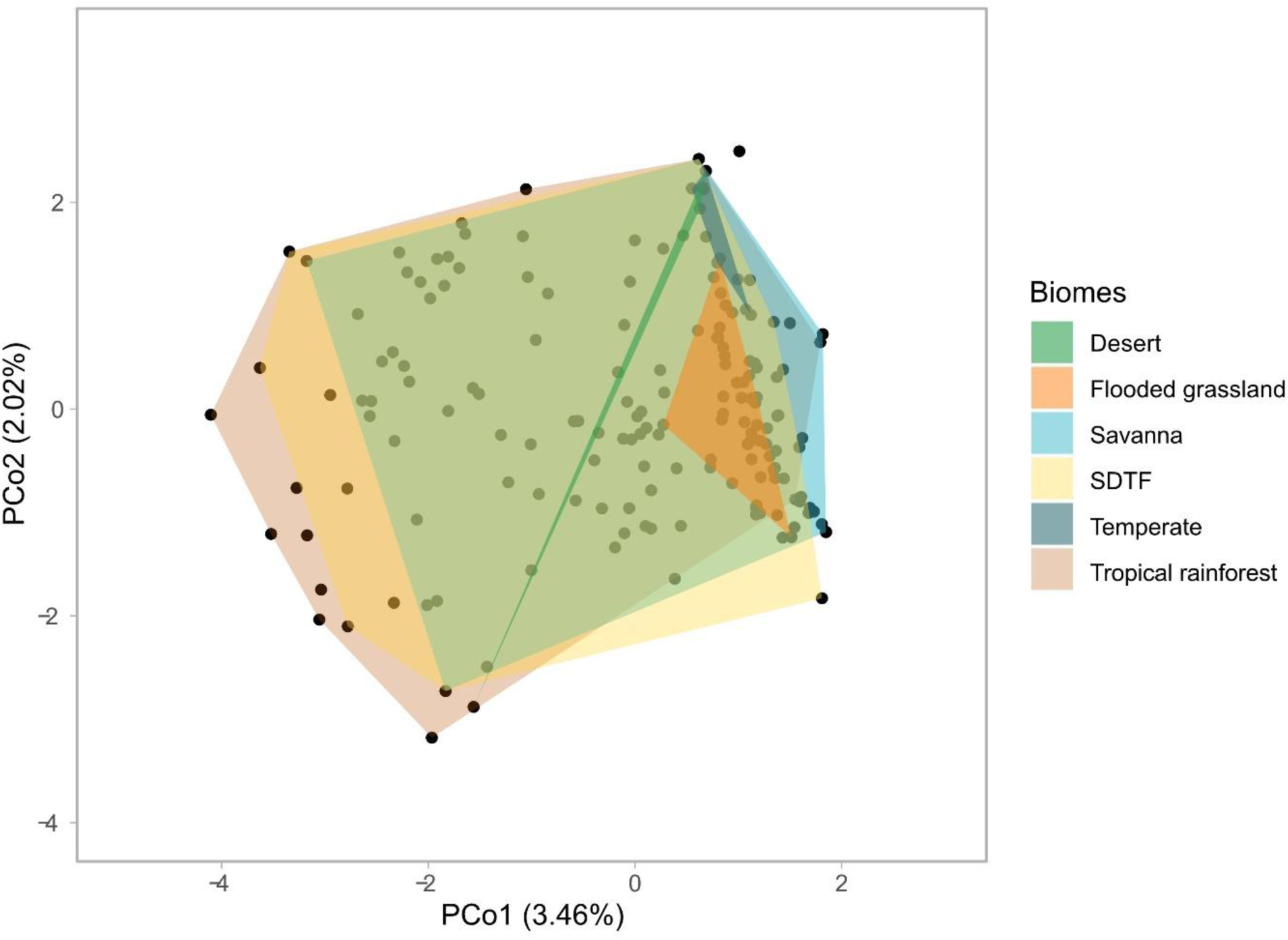
Morphospace of Mimosa and Stryphnodendron clades with polygons indicating the convex hull for each ecological group. Values in parentheses indicate the percentage of variance explained by each axis.

Disparity metrics reinforce what is observed in the morphospace, indicating a greater disparity in pollen morphology of tropical rainforest and SDTF taxa, while savanna stands as an intermediary case, considering both morphospace extent (Fig. 3C) and density (Supplementary Table S3). Although the other biomes present lesser disparity, desert shows scattered morphospace occupancy (Supplementary Table S3). We found that the medians of morphospace extent values differ significantly among all biomes, but there is also a high probability of overlap between flooded grassland and temperate, as well as between tropical rainforest and SDTF.

Despite limited pollen data allowing for the analysis of only five out of the twelve delimited ecological lineages, there is notable variation in disparity among lineages occurring in SDTF (Fig. 3D). Three independent SDTF lineages were analyzed, two from the Mimosa clade and one from the Stryphnodendron clade. One of Mimosa clade’s SDTF lineages, here addressed as “Mimosa SDTF 1”, shows greater disparity than the other two lineages from the same biome (Fig. 3D, Supplementary Table S3) that probably overlap in extent values (Fig. 4C). Pollen disparity of the Mimosa SDTF 1 lineage is in fact more comparable to those of savanna and tropical rainforest, with extent values probably overlapping with those of the Stryphnodendron clade tropical rainforest lineage.

## DISCUSSION

Here, we used disparity analyses to investigate the pollen morphological diversity of different ecological groups and lineages in the Mimosa and Stryphnodendron clades of mimosoid legumes. Overall, two genera of the Mimosa clade, *Adenopodia* and *Piptadenia*, share similar patterns of pollen disparity with members of the Stryphnodendron clade, while the genus *Mimosa* occupies a distinct area of the pollen morphospace (Figs. 2, 3B).

As for ecological groups, tropical rainforest and SDTF have the highest median disparity, considering both morphospace extent (Fig. 3C) and density (Table S3). In general, there is a low probability of overlap in values of morphospace extent for ecological groups, but exceptions occur between temperate and flooded grassland, as well as SDTF and tropical rainforest (Fig. 4). Finally, not all independent lineages of SDTF share similar disparity. Instead, although they are not necessarily placed in the same region, an SDTF lineage of the Mimosa clade and a tropical rainforest lineage of the Stryphnodendron clade occupy similar ranges of the morphospace (Fig. 4).

### Pollen disparity in Mimosa and Stryphnodendron clades

Distribution of lineages in the phylomorphospace and disparity metrics (Figs. 2, 3) indicate distinct patterns of pollen evolution across members of the Mimosa and Stryphnodendron clades. While *Mimosa* is placed in its own area of the morphospace, all remaining genera share the same region, which is mainly characterized by larger dispersal units with 8 to 16 grains. Nonetheless, the overall distribution of species in the morphospace and the lack of overlap between some genera shows that these lineages are not in evolutionary stasis. However, similarly to other traits and organisms (e.g., Santos *et al*., 2019), pollen evolution in *Adenopodia*, *Piptadenia* and the Stryphnodendron clade seems to be limited to particular morphologies, which are recurrently accessed across time.

This pattern suggests the existence of distinct evolutionary regimes among lineages of the Mimosa and Stryphnodendron clades. One regime, likely ancestral, would apply for the Stryphnodendron clade, *Adenopodia* and *Piptadenia*, and another for *Mimosa*. Although we don’t test this idea here, additional evidence lends support to it. First, while members of the Stryphnodendron clade, *Adenopodia* and *Piptadenia* occur primarily in tropical rainforests, *Mimosa* is particularly diverse in drier biomes (e.g., Savanna and SDTF) (Borges, Simon, Morales, *et al*., 2024; Borges, Simon, Ribeiro, *et al*., 2024) and includes a number of biome shifts (Ringelberg et al. 2023), indicating it may be more evolutionarily labile. Second, and most importantly, pollen development in *Mimosa* differs from other genera in the Mimosa and Stryphnodendron clades (Seijo and Solis Neffa, 2004; Capucho and Teixeira, 2020) and could be related to a release of biological constraints applying to the other taxa studied here. Third, overlap in morphospace extent is much more likely among genera of the Stryphnodendron clade, *Adenopodia* and *Piptadenia* (Fig. 4A). Such a link between changes in pollen evolutionary regimes and the overcoming of biological constraints has been made before for Myrtales pollen (Kriebel *et al*., 2017) and may help to explain the different placement of *Mimosa* in the morphospace and its highest pollen disparity. Nonetheless, although *Mimosa* has the highest values for sum of variances, *Piptadenia* and some lineages of the Stryphnodendron clade, such as *Gwilymia* and *Pityrocarpa*, also stand with high disparity values when compared to the remaining genera.

Although the high diversity of *Mimosa* pollen has been highlighted before (Caccavari, 2002), we showed that pollen evolution in the genus involved a shift to a novel area of the morphospace with no regressions to the ancestral-like morphologies seen in other taxa of the Mimosa and Stryphnodendron clades. As hypothesized by Barneby (1991), this novel area of the morphospace comprises smaller dispersal units with fewer pollen grains (mostly tetrads). These innovations may have enabled a burst in morphological evolution, increasing pollen disparity (Lupia, 1999). It is possible this expansion of pollen disparity in *Mimosa* could be linked to variations in diversification rates inferred for the genus (Koenen *et al*., 2013; Vasconcelos *et al*., 2020; Ringelberg *et al*., 2023). However, shifts in diversification rates do not seem to match shifts in the number of grains of the dispersal unit across the *Mimosa* phylogeny (Santos-Silva *et al*., 2013), for example. Thus, changes in the diversification of pollen morphology may have other origins for these plants.

As hinted above, novel pollen forms in *Mimosa* could be related to changes in development and mode of reproduction. The few ontogenetic studies available have shown that *Mimosa* differs from other closely related mimosoids not just in the number of aggregated cells (i.e., tetrads instead of polyads), but also in pollen sac morphology and the number of microspore mother-cells (Seijo and Solis Neffa, 2004; Capucho and Teixeira, 2020). *Mimosa* also seems to produce higher numbers of pollen grains per anther in several species (Seijo, personal observation). High pollen yield is not common in mimosoids, in which the relationship between the number of pollen grains and the number of ovules (P/O ratio) in each flower is usually low (Kenrick and Knox, 1982) and associated with high efficiency in pollen dispersal (Capucho and Teixeira, 2020). Thus, the new area of the pollen morphospace occupied by *Mimosa* (Fig. 2) is clearly linked to developmental changes. These changes, in turn, may have occurred in an adaptive context, such as modifications in pollination regime.

Different pollination features are at the center of hypotheses to explain the evolution of aggregated dispersal units. One hypothesis postulates that aggregated units of the Mimoseae lower the occurrence of mixed loads of self and cross-pollen in the stigma, a shortcoming that can lead to ovule abortion in some species (Wyatt and Lipow, 2021). *Acacia* Mill. and *Inga*, for example, produce large, many-grained polyads that fill the entire stigmatic cavity, blocking the entrance of additional polyads (Seijo and Solis Neffa, 2004). In *Mimosa*, the stigmas of *M. bimucronata* and *M. caesalpiniifolia* fit a single polyad, also corroborating this hypothesis, but *M. microphylla* deviates from this pattern, with stigmas that fit up to six tetrads (Wyatt and Lipow, 2021; Guglielmi and Teixeira, 2024). Moreover, the reversal towards lower numbers of pollen grains in each dispersal unit seen for *Mimosa* (Fig. 2) actually goes against Wyatt and Lipow (2021) hypothesis. Thus, the *Mimosa* morphological shift would fit better a scenario where aggregation in many-grained polyads is maladaptive in some cases (Harder and Johnson, 2008).

In this context, another hypothesis places aggregated units in a trade-off framework, enhancing pollen transport efficiency in some cases, such as infrequent pollinators, but reducing efficiency in others, such as frequent self-pollination or bee grooming (Harder and Johnson, 2008). However, pollinator diversity and abundance vary with elevation due to thermal tolerances and resource availability (Lara-Romero *et al*., 2019). For example, in a montane site in Brazil, the abundance of bees, a main mimosoid pollinator (Vogel *et al*., 2005; Banks and Rudall, 2016), declines at higher elevations (dos Santos *et al*., 2020). Yet, sites of high-elevation are the main centers of *Mimosa* diversity and endemism (Simon and Proença, 2000). In this context, the evolutionary change in the pollen morphology of the genus could not be explained by increased pollinator frequency, as discussed above (Harder and Johnson, 2008). Because of that, we propose that the shift from larger polyads to smaller tetrads might have happened as an adaptation to factors reducing the chance of pollen reaching conspecific stigmas, such as self-pollination or bee grooming (Harder and Johnson, 2008). Thus, instead of *Mimosa* pollen grains co-evolving with stigmas to avoid abortion from mixed pollen loads (Wyatt and Lipow, 2021), their morphology may have shifted from polyads to tetrads to maximize pollen transfer efficiency (Harder and Johnson, 2008), even at the cost of mixed pollen loads in the stigma.

### Pollen disparity across different biomes

In addition to phylogenetic relationships and biotic interactions, abiotic factors, such as temperature and humidity, may also influence pollen form (Ejsmond *et al*., 2011; Luo *et al*., 2015; Kriebel *et al*., 2017). These abiotic factors are particularly important in the evolution of traits modulating changes in pollen shape in response to water loss at the final stages of maturation (harmomegathy), such as exine thickness and ornamentation, grain size (long and short diameters), and number or type of apertures (van der Ham and van Heuven, 1989; Franchi *et al*., 2002; Katifori *et al*., 2010; Yu *et al*., 2018). Although variations in some of these traits are seen in the Mimosa and Stryphnodendron clades (Supplementary Table S5), our results suggest that pollen disparity is not limited by abiotic conditions of different biomes. Indeed, even though some biomes show distinct pollen disparity (e.g., Tropical rainforest versus temperate; Fig. 4B), biomes of different nature may share similar levels of morphological diversity (e.g., Tropical rainforest and SDTF; Fig. 5).

Few studies have investigated the influence of different habitat conditions on pollen morphology, with the majority focused on intraspecific variation (Li *et al*., 2011; Fatmi *et al*., 2020; Wrońska-Pilarek *et al*., 2023). Despite the difference in evolutionary scale, these studies at the species level bear similarities with our findings in a macroevolutionary context. For example, although pollen morphology of *Convallaria majalis* L. (Asparagaceae) is somewhat correlated with habitat variation, similar characters occur in markedly different habitats (Wrońska-Pilarek *et al*., 2023). In most cases, the relationship between morphology and habitat occurs in some of the harmomegathy-related characters highlighted above. In *C. majalis* again, among all the pollen characters investigated (e.g., ornamentation, outline, diameters, exine thickness), only exine thickness was strongly correlated to habitat (Wrońska-Pilarek *et al*., 2023). Similarly, only variation in size and shape of *Atriplex halimus* L. (Amaranthaceae) pollen is related to abiotic factors (Fatmi *et al*., 2020). As we used a multivariate approach here, it is possible that variations in individual traits of tetrads or polyads could be correlated to specific environmental variables, particularly as these aggregated units also possess harmomegathic adaptations (Guinet, 1986; Franchi *et al*., 2002). Nonetheless, our results indicate that the overall morphology of aggregated dispersal units is decoupled from environmental factors.

Despite a lack of clear separation in the morphospace, we found that pollen disparity varies among ecological groups. As expected from different species richness (Guinet and Caccavari, 1992; Caccavari, 2002), tropical rainforest and SDTF show higher disparities, while biomes where the taxonomic diversity of the Mimosa and Stryphnodendron clades is limited, such as temperate and flooded grasslands, are less morphologically diverse. Nonetheless, as indicated by values of the Bhattacharyya coefficient, there is a high chance of overlap between the disparity of some ecological groups. In particular, tropical rainforest and SDTF, two biomes with very different climatic conditions (Pennington and Lavin, 2016), show very similar values of morphospace occupation (Fig. 3C, Fig. 4), meaning that lineages occurring in both biomes are equally able to explore the pollen morphospace, reaching similar levels of disparity. Even the relatively wide observable disparity seen for desert, which includes a low diversity of taxa, exemplifies how harsh conditions do not constrain pollen morphology in these plants. Again, our results suggest that environmental factors did not strongly shape pollen disparity in the Mimosa and Stryphnodendron clades.

Results for ecological lineages reinforce this hypothesis, as independent SDTF lineages do not show convergent disparity. This suggests that the dry environment of the SDTFs, a potential restrictor of pollen morphology (Yu *et al*., 2018), does not limit the disparity of tetrads and polyads within the Mimosa and Stryphnodendron clades. On the other hand, the differences in disparity seen between these independent SDTF lineages are likely more related to historical factors, such as developmental constraints, for example (Oyston *et al*., 2015; Jablonski, 2020).

It is important to note that some of the initially delimited ecological lineages, such as the savanna lineages in the genus *Stryphnodendron*, and the *Mimosa* lineages from tropical rainforests (Supplementary Fig. S2), could not be analyzed due to lack of pollen morphological data. This problem stems in part from the trade-off of choosing biomes to delimit ecological lineages (Casali *et al*., 2023). One potential solution would be to classify lineages into broader ecological patterns, such as precipitation or temperature regimes. Precipitation, for example, has been shown to influence species turnover across space (Neves *et al*., 2020; Ringelberg *et al*., 2023) and could be correlated to pollen disparity. Nevertheless, by focusing on biomes, we use categories that are connected to major evolutionary trends operating across the phylogeny of land plants (e.g., Jaramillo, 2023; Hagelstam-Renshaw *et al*., 2024). Regardless of analytical choices, it is still important to expand palynological studies to fill gaps in the description of pollen for most taxa in the Mimosa and Stryphnodendron clades, which would, in turn, support macroevolutionary studies such as this.

Altogether, the evidence presented here suggests that overall evolution of pollen dispersal units in the Mimosa and Stryphnodendron clades is primarily related to factors other than ecology, such as evolutionary history and development. This differs at least in part from what has been shown for Myrtales, in which pollen disparity is correlated with latitude, likely as a proxy for harmomegathy (Kriebel *et al*., 2017). Thus, the lack of environmentally driven variation in the Mimosa and Stryphnodendron clades is likely an example of phylogenetic conservation of pollen morphology (Yu *et al*., 2019), which is commonly observed in different angiosperm groups (e.g., Lu *et al*., 2015; Luo *et al*., 2015).

We found that the Mimosa and Stryphnodendron clades exhibit dissimilar patterns of disparity, with the Mimosa clade showing greater disparity, primarily due to the shift of the genus *Mimosa* to a novel area of the pollen morphospace. Our findings suggest that *Mimosa* pollen dispersal units overcame constraints, most likely developmental, or followed a different evolutionary regime, probably linked to increased pollen transfer efficiency to stigmas of conspecific individuals. Furthermore, by integrating pollen morphology and ecological data we highlighted tropical rainforest and SDTF taxa, biomes with very different climatic conditions, as the most disparate ecological groups, and showed their disparity overlaps. Additionally, we have also found that distinct lineages occurring in the dry environments of SDTFs do not converge in pollen morphology. However, the similarities in disparity found between very different biomes (SDTF and Tropical rainforest), coupled with the absence of such a pattern between SDTF lineages, suggest that the evolution of aggregate dispersal units is largely decoupled from ecology. Thus, although ecological factors may have played a role in pollen disparification within particular lineages of the Mimosa and Stryphnodendron clades, the overall consistent pattern of morphological conservatism found in pollen grains (Traverse, 2007; Kriebel *et al*., 2017; Jardine *et al*., 2022) is not blurred by ecology. Our findings emphasize the idiosyncratic nature of pollen disparification across plant lineages and highlight the importance of considering both extrinsic and intrinsic components when investigating disparity-related questions. However, here we show that the morphological evolution of aggregated dispersal units is mainly driven by lineage-specific history, not by ecology.

## ACKNOWLEDGEMENTS

We thank Monique Maianne and Yago Barros-Souza for their support and useful comments at different stages of this research.

## DATA AVAILABILITY

All original and legacy data, as well as code used for analyses will be provided on the peer-reviewed publication.

