## Supplementary information for "Morphological innovation and lineage-specific history drive disparification in the aggregated pollen of mimosoid plants"

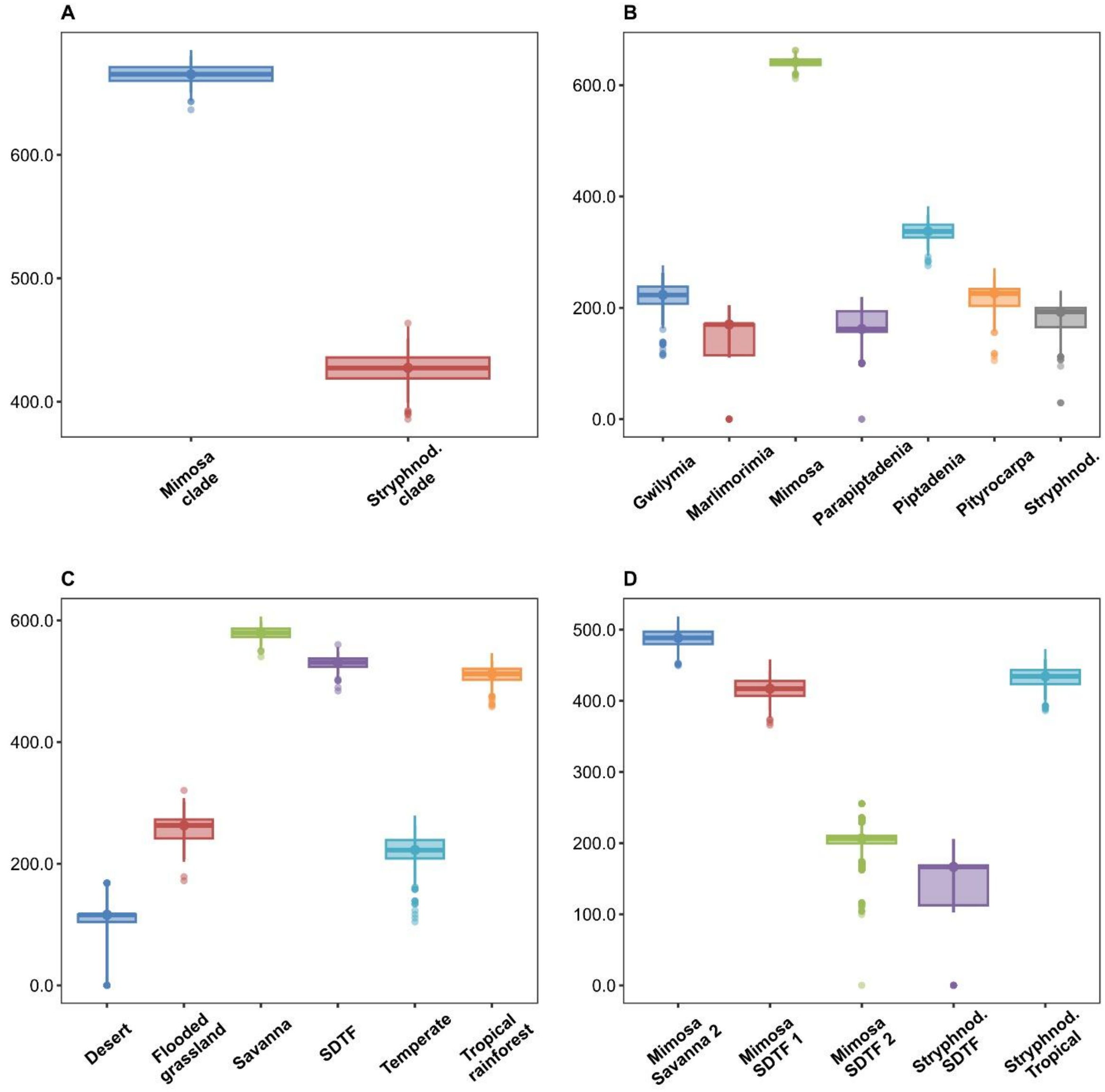


Fig. S1. Morphological disparity (sum of ranges) for groups. Comparisons between Mimosa and Stryphnodendron clades (A), within-clades genera (B), ecological groups (C) and ecological lineages (D). Boxplots show the distribution of values from 1000 bootstrap replicates.

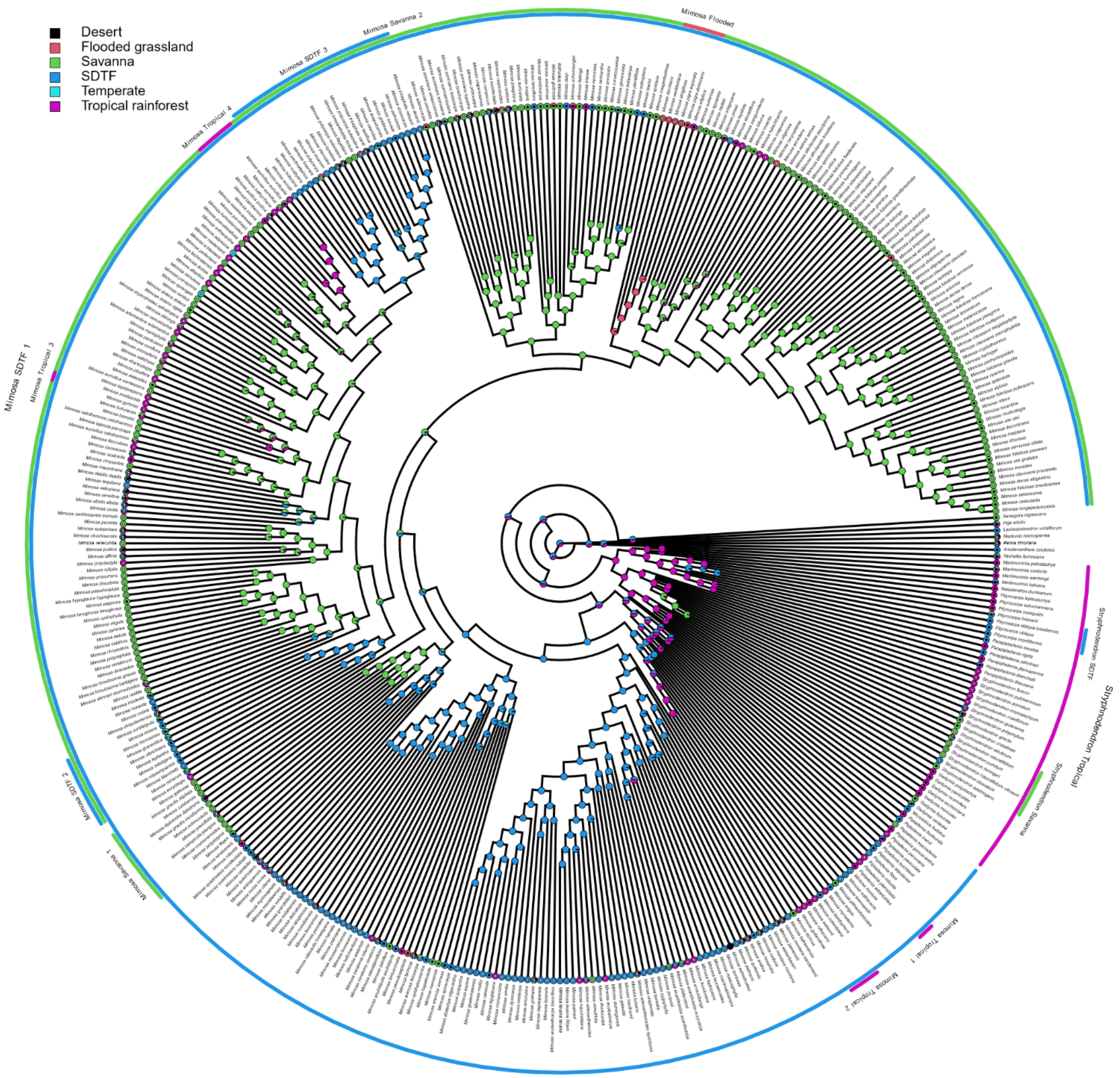


Fig. S2. Ancestral biomes estimate for Mimosa and Stryphnodendron clades, with delimitated lineages represented around the phylogeny. Species with and without pollen data available have black and white dots inside tips, respectively. Lineages with sufficient pollen data for analysis were Mimosa Savanna 2, Mimosa SDTF 1, Mimosa SDTF 2, Stryphnodendron SDTF and Stryphnodendron tropical.

Table S1. *Morphospace occupancy of lineages and ecological groups calculated from the sum of ranges (SR) metric.*

| Subdivision | Group | Sample_size | SR_Obs | Median_BS | CI (BS) |
|---|---|---|---|---|---|
| Genera | Gwilymia | 7 | 276.5 | 223.2 | 168.4-262.9 |
| Genera | Marlimorimia | 4 | 204.9 | 170 | 110.4-204.9 |
| Genera | Mimosa | 135 | 704.5 | 641.3 | 625.6-657.4 |
| Genera | Parapiptadenia | 5 | 219.3 | 161.9 | 101.4-219.3 |
| Genera | Piptadenia | 18 | 397.9 | 337.4 | 302.9-366.9 |
| Genera | Pityrocarpa | 7 | 271.1 | 226 | 163.4-255.7 |
| Genera | Stryphnodendron | 6 | 230.8 | 192.7 | 117.4-221 |
| Clades | Mimosa clade | 155 | 729 | 665.3 | 650.5-680.1 |
| Clades | Stryphnodendron clade | 32 | 489.1 | 427.4 | 399.2-451.2 |
| Biomes | Desert | 3 | 168.1 | 116 | 0-168.1 |
| Biomes | Flooded grassland | 9 | 320.5 | 263 | 207.4-301.9 |
| Biomes | Savanna | 83 | 646.1 | 579.7 | 558.2-598.6 |
| Biomes | SDTF | 64 | 594.3 | 530.8 | 510.8-549.1 |
| Biomes | Temperate | 7 | 279 | 222.6 | 167.7-262.6 |
| Biomes | Tropical rainforest | 61 | 576.9 | 512.1 | 482.7-533.9 |
| Ecological lineages | Mimosa_Savanna_2 | 46 | 555.7 | 488.7 | 462.7-510.7 |
| Ecological lineages | Mimosa_SDTF_1 | 33 | 476.5 | 417.1 | 387.1-441.3 |
| Ecological lineages | Mimosa_SDTF_2 | 6 | 255.3 | 206.6 | 162.4-235.7 |
| Ecological lineages | Stryphnodendron SDTF | 4 | 205.8 | 166.5 | 102.6-205.8 |
| Ecological lineages | Stryphnodendron Tropical | 36 | 495 | 434.5 | 401.4-458.3 |

Median and confidence interval (CI) values for bootstrapped and rarefied data (BS).

Table S2. *Morphospace occupancy of lineages and ecological groups calculated from the sum of variances (SV) metric.*

| Subdivision | Group | Sample size | SV (Obs.) | Median (BS) | CI (BS) |
|---|---|---|---|---|---|
| Genera | Gwilymia | 7 | 64.16 | 55.58 | 43.42-61.39 |
| Genera | Marlimorimia | 4 | 63.92 | 53.14 | 30.53-63.92 |
| Genera | Mimosa | 135 | 65.01 | 64.52 | 64.07-64.99 |
| Genera | Parapiptadenia | 5 | 63 | 53.2 | 25.84-63 |
| Genera | Piptadenia | 18 | 65.14 | 61.66 | 58.39-63.62 |
| Genera | Pityrocarpa | 7 | 63.65 | 55.77 | 42.73-61.23 |
| Genera | Stryphnodendron | 6 | 61.43 | 53.35 | 36.92-57.84 |
| Clades | Mimosa clade | 155 | 66.07 | 65.64 | 65.04-66.14 |
| Clades | Stryphnodendron clade | 32 | 64.41 | 62.5 | 60.73-63.79 |
| Biomes | Desert | 3 | 69.34 | 49.27 | 0-69.34 |
| Biomes | Flooded grassland | 9 | 64.19 | 57.62 | 50.03-62.21 |
| Biomes | Savanna | 83 | 64.95 | 64.16 | 63.36-64.91 |
| Biomes | SDTF | 64 | 66.9 | 65.87 | 65.05-66.54 |
| Biomes | Temperate | 7 | 62.88 | 54.6 | 42.01-60.38 |
| Biomes | Tropical rainforest | 61 | 67.39 | 66.34 | 65.48-66.99 |
| Ecological lineages | Mimosa_Savanna_2 | 46 | 63.11 | 61.79 | 60.71-62.57 |
| Ecological lineages | Mimosa_SDTF_1 | 33 | 65.84 | 63.89 | 62.4-64.91 |
| Ecological lineages | Mimosa_SDTF_2 | 6 | 63.94 | 55.16 | 38.27-60.45 |
| Ecological lineages | Stryphnodendron SDTF | 4 | 63.78 | 51.63 | 28.02-63.78 |
| Ecological lineages | Stryphnodendron Tropical | 36 | 66.87 | 65.09 | 63.22-66.29 |

Median and confidence interval (CI) values for bootstrapped and rarefied data (BS).

Table S3. *Morphospace occupancy of lineages and ecological groups calculated from the mean pairwise distance (MPD) metric.*

| Subdivision | Group | MPD |
|---|---|---|
| Genera | Gwilymia | 0.74 |
| Genera | Marlimorimia | 0.72 |
| Genera | Mimosa | 0.81 |
| Genera | Parapiptadenia | 0.63 |
| Genera | Piptadenia | 0.82 |
| Genera | Pityrocarpa | 0.69 |
| Genera | Stryphnodendron | 0.5 |
| Clades | Mimosa_clade | 0.9 |
| Clades | Stryphnodendron_clade | 0.76 |
| Biomes | Desert | 1.18 |
| Biomes | Flooded_grassland | 0.74 |
| Biomes | Savanna | 0.81 |
| Biomes | SDTF | 0.98 |
| Biomes | Temperate | 0.63 |
| Biomes | Tropical_rainforest | 1.02 |
| Ecological lineages | Mimosa_Savanna_2 | 0.65 |
| Ecological lineages | Mimosa_SDTF_1 | 0.89 |
| Ecological lineages | Mimosa_SDTF_2 | 0.72 |
| Ecological lineages | Stryphnod_SDTF | 0.7 |
| Ecological lineages | Stryphnod_Tropical | 0.97 |

Table S4. *Wilcoxon p-test and Bhattacharyya Coefficient (BC) for group's bootstrapped sum of variances (SR) values.*

| Subdivision | Groups | Statistic W | P-value | BC |
|---|---|---|---|---|
| Genera | Gwilymia : Marlimorimia | 735118 | >0.05 | 0.34 |
| Genera | Gwilymia : Mimosa | 128 | >0.05 | 0.04 |
| Genera | Gwilymia : Parapiptadenia | 678139 | >0.05 | 0.74 |
| Genera | Gwilymia : Piptadenia | 52999 | >0.05 | 0.37 |
| Genera | Gwilymia : Pityrocarpa | 519064 | 0.28 | 0.92 |
| Genera | Gwilymia : Stryphnodendron | 709467 | >0.05 | 0.64 |
| Genera | Marlimorimia : Mimosa | 504 | >0.05 | 0.29 |
| Genera | Marlimorimia : Parapiptadenia | 445089 | >0.05 | 0.57 |
| Genera | Marlimorimia : Piptadenia | 83601 | >0.05 | 0.09 |
| Genera | Marlimorimia : Pityrocarpa | 276773 | >0.05 | 0.46 |
| Genera | Marlimorimia : Stryphnodendron | 431900 | >0.05 | 0.49 |
| Genera | Mimosa : Parapiptadenia | 1000000 | >0.05 | 0 |
| Genera | Mimosa : Piptadenia | 999281 | >0.05 | 0.07 |
| Genera | Mimosa : Pityrocarpa | 1000000 | >0.05 | 0.08 |
| Genera | Mimosa : Stryphnodendron | 1000000 | >0.05 | 0 |
| Genera | Parapiptadenia : Piptadenia | 51497 | >0.05 | 0.26 |
| Genera | Parapiptadenia : Pityrocarpa | 333620 | >0.05 | 0.77 |
| Genera | Parapiptadenia : Stryphnodendron | 485264 | 0.28 | 0.77 |
| Genera | Piptadenia : Pityrocarpa | 952568 | >0.05 | 0.41 |
| Genera | Piptadenia : Stryphnodendron | 993059 | >0.05 | 0.17 |
| Genera | Pityrocarpa : Stryphnodendron | 687572 | >0.05 | 0.67 |
| Clades | Mimosa_clade : Stryphnodendron_clade | 1000000 | >0.05 | 0 |
| Biomes | Desert : Flooded_grassland | 230334 | >0.05 | 0.09 |
| Biomes | Desert : Savanna | 222000 | >0.05 | 0 |
| Biomes | Desert : SDTF | 222000 | >0.05 | 0 |
| Biomes | Desert : Temperate | 285137 | >0.05 | 0.16 |
| Biomes | Desert : Tropical_rainforest | 222000 | >0.05 | 0 |
| Biomes | Flooded_grassland : Savanna | 525 | >0.05 | 0.04 |
| Biomes | Flooded_grassland : SDTF | 0 | >0.05 | 0 |
| Biomes | Flooded_grassland : Temperate | 717277 | >0.05 | 0.84 |
| Biomes | Flooded_grassland : Tropical_rainforest | 0 | >0.05 | 0 |
| Biomes | Savanna : SDTF | 1177 | >0.05 | 0.08 |
| Biomes | Savanna : Temperate | 1000000 | >0.05 | 0.01 |
| Biomes | Savanna : Tropical_rainforest | 153 | >0.05 | 0.03 |
| Biomes | SDTF : Temperate | 1000000 | >0.05 | 0 |
| Biomes | SDTF : Tropical_rainforest | 189827 | >0.05 | 0.8 |
| Biomes | Temperate : Tropical_rainforest | 0 | >0.05 | 0 |
| Ecological lineages | Mimosa_Savanna_2 : Mimosa_SDTF_1 | 8471 | >0.05 | 0.19 |
| Ecological lineages | Mimosa_Savanna_2 : Mimosa_SDTF_2 | 983550 | >0.05 | 0.04 |

| Subdivision | Groups | Statistic W | P-value | BC |
|---|---|---|---|---|
| Ecological lineages | Mimosa_Savanna_2 : Stryphnod_SDTF | 905000 | >0.05 | 0 |
| Ecological lineages | Mimosa_Savanna_2 : Stryphnod_Tropical | 596 | >0.05 | 0.06 |
| Ecological lineages | Mimosa_SDTF_1 : Mimosa_SDTF_2 | 991360 | >0.05 | 0.1 |
| Ecological lineages | Mimosa_SDTF_1 : Stryphnod_SDTF | 959055 | >0.05 | 0.28 |
| Ecological lineages | Mimosa_SDTF_1 : Stryphnod_Tropical | 119046 | >0.05 | 0.65 |
| Ecological lineages | Mimosa_SDTF_2 : Stryphnod_SDTF | 675247.5 | >0.05 | 0.56 |
| Ecological lineages | Mimosa_SDTF_2 : Stryphnod_Tropical | 1504 | >0.05 | 0.07 |
| Ecological lineages | Stryphnod_SDTF : Stryphnod_Tropical | 6555 | >0.05 | 0.23 |

Table S5. *Pollen data from taxa of the Mimosa and Stryphnodendron clades. Character state names can be found in Table 1 of the original paper.*

| Taxon names | Literature source | D.U. Number of grains | D.U. Outline | Pollen arrangement | Pollen ornamentation | D.U. Shorter diameter | D.U. Longer diameter | Pollen exine thickness |
|---|---|---|---|---|---|---|---|---|
| *Adenopodia gymnantha* | Caccavari 2002 | 5 | 1 | | 7 | | 28.00 | |
| *Adenopodia oaxacana* | Caccavari 2002 | 5 | 1 | | 7 | | 31.00 | |
| *Anadenanthera colubrina* | Cruz et al. 2018; Ribeiro et al. 2018 | 5 | 0&1 | 2 | 0&3 | 39.00 | 38.80 | 2.00 |
| *Anadenanthera colubrina var. cebil* | Caccavari 2002; Ribeiro et al. 2018 | 4 | 1&2 | 2 | 0&6 | 35.07 | 37.78 | 2.00 |
| *Anadenanthera colubrina var. colubrina* | Buril et al. 2010 | 5 | | | 0 | 33.90 | 37.20 | 2.50 |
| *Anadenanthera peregrina* | Ribeiro et al. 2018; Caccavari 2002 | 5 | 1 | 2 | 0 | 29.77 | 34.55 | 1.00 |
| *Gwilymia coriacea* | Barduzzi et al. 2024; Guinet & Caccavari 1992 | 5 | 1&2 | | 6 | 30.80 | 34.10 | 1.00 |
| *Gwilymia fissurata* | Caccavari 2002; Guinet & Caccavari 1992 | 5 | 2 | 5 | 6 | 40.50 | 63.30 | 1.80 |
| *Gwilymia moricolor* | Guinet & Caccavari 1992 | 5 | 1&2 | 5 | 0&1 | 44.50 | 57.60 | 1.75 |
| *Gwilymia occhioniana* | Guinet & Caccavari 1992 | 3 | 1&2 | 1 | 6 | 31.10 | 47.90 | 1.70 |
| *Gwilymia paniculata* | Guinet & Caccavari 1992 | 5 | 2 | 3 | 0&1 | | 44.50 | 1.75 |
| *Gwilymia polystachya* | Guinet & Caccavari 1992 | 3 | 1&2 | 1 | 6 | 34.10 | 46.60 | 1.70 |
| *Gwilymia racemifera* | Guinet & Caccavari 1992 | 5 | 1&2 | 0&5 | 0&5 | 34.10 | 55.00 | 1.75 |
| *Inga edulis* | Braga et al. 2012 | 6 | 1 | | 6 | 50.00 | 100.00 | 1.70 |
| *Lachesiodendron viridiflorum* | Barduzzi et al. 2024 | 3 | | 5 | 6 | 24.07 | 35.74 | 0.85 |
| *Marlimorimia bahiana* | Caccavari 2002; Ribeiro et al. 2018 | 5 | 2 | 0&3 | 4 | 32.70 | 40.58 | 1.00 |
| *Marlimorimia contorta* | Barduzzi et al. 2024; Caccavari 2002; Ribeiro et al. 2018 | 5 | 2 | 0&4&5 | 4 | 27.51 | 37.52 | 0.61 |
| *Marlimorimia psilostachya* | Caccavari 2002 | 5 | 2 | 3 | 0 | | 40.00 | |
| *Marlimorimia warmingii* | Barth & Yoneshigue 1966 | 4&5 | 1&2 | 1 | 4 | 37.80 | 46.25 | 2.50 |
| *Microlobius foetidus* | Barduzzi et al. 2024; Caccavari 2002 | 3 | 2 | 5 | 7 | 17.28 | 24.68 | 0.22 |
| *Mimosa acantholoba* | Liau-Kang et al. 2024 | 3 | 1 | 3 | 0 | 8.53 | 12.80 | 0.21 |
| *Mimosa acutistipula* | Santos-Silva et al. 2013 | 3 | 1 | 3 | 6 | 12.10 | 17.90 | |
| *Mimosa acutistipula var. acutistipula* | Lima et al. 2008 | 3 | | 3&5 | 0&6 | 12.60 | 16.90 | 1.00 |
| *Mimosa adenocarpa* | Liau-Kang et al. 2024 | 2 | 0 | 0&5 | 0&6 | 11.07 | 14.47 | 0.44 |
| *Mimosa adenophylla* | Santos-Silva et al. 2013 | 2 | 1 | 0&5 | 7 | 15.50 | 22.00 | |
| *Mimosa adenophylla var. armandiana* | Lima et al. 2008 | 1&2 | | 0&3 | 0&6 | 16.40 | 21.50 | 1.00 |
| *Mimosa adenophylla var. mitis* | Lima et al. 2008 | 1&2 | | 0&3 | 0&6 | 15.40 | 22.00 | 1.00 |
| *Mimosa adenotricha* | Liau-Kang et al. 2024 | 2 | 1 | 0&4&5 | 0&6 | 11.94 | 16.34 | 0.34 |
| *Mimosa aguapeia* | Liau-Kang et al. 2024 | 2 | 1 | 3 | 0 | 18.50 | 24.80 | |
| *Mimosa albolanata var. brasiliana* | Liau-Kang et al. 2024 | 2 | 1 | 0&3 | 0 | 13.10 | 20.40 | |
| *Mimosa amnis-atri* | Santos-Silva et al. 2013 | 2 | 1 | 3 | 2&3 | 13.10 | 12.70 | |

| Taxon names | Literature source | D.U. Number of grains | D.U. Outline | Pollen arrangement | Pollen ornamentation | D.U. Shorter diameter | D.U. Longer diameter | Pollen exine thickness |
|---|---|---|---|---|---|---|---|---|
| *Mimosa apodocarpa* | Santos-Silva et al. 2013 | 2 | 1 | 3 | 0&6 | 14.50 | 14.20 | |
| *Mimosa arenosa* | Buril et al. 2010 | 3 | 1&2 | 3&4 | 0&6 | 10.10 | 13.80 | |
| *Mimosa arenosa var. arenosa* | Lima et al. 2008 | 3 | 1 | 3 | 0&6 | 9.70 | 14.20 | 1.20 |
| *Mimosa artemisiana* | Santos-Silva et al. 2013 | 2 | 1 | | 3 | 15.50 | 25.00 | |
| *Mimosa aurivillus var. aurivillus* | Liau-Kang et al. 2024 | 2 | 0 | 0&5 | 6 | 11.28 | 14.00 | 0.83 |
| *Mimosa bimucronata* | Cruz et al. 2018; Liau-Kang et al. 2024; Barth & Yoneshigue 1966; Barth 1973 | 3 | 1 | 0&5 | 3&4&6 | 10.86 | 14.86 | 0.17 |
| *Mimosa bimucronata var. bimucronata* | Lima et al. 2008 | 3 | 1 | 0 | 0&6 | 11.20 | 15.40 | 1.00 |
| *Mimosa borboremae* | Lima et al. 2008 | 2 | | 0&5 | 0&6 | 8.70 | 11.90 | 0.50 |
| *Mimosa caerulea* | Medina-Acosta-2018 | 2 | 0&1 | 5 | 2&3 | | 14.00 | 1.06 |
| *Mimosa caesalpiniifolia* | Lima et al. 2008 | 3 | 1 | 3 | 0&6 | 9.80 | 12.90 | 1.00 |
| *Mimosa calcicola* | Medina-Acosta-2018 | 3 | 1 | 3&4 | 2&3 | 12.88 | 16.73 | 0.90 |
| *Mimosa campicola var. campicola* | Lima et al. 2008 | 2 | | 0&3 | 0&6 | 23.70 | 29.50 | 1.00 |
| *Mimosa candollei* | Liau-Kang et al. 2024; Buril et al. 2010; Lima et al. 2008 | 2 | 1&2 | 3&5 | 0&6 | 29.37 | 35.67 | 0.30 |
| *Mimosa capito* | Liau-Kang et al. 2024 | 2 | 1 | 0 | 0 | 12.70 | 20.10 | |
| *Mimosa ceratonia* | Cruz et al. 2018 | 5 | 1 | 4 | 3 | 20.60 | 23.50 | 2.00 |
| *Mimosa ceratonia var. pseudo-obovata* | Liau-Kang et al. 2024 | 2&3 | 1 | 5 | 0&3 | 10.84 | 14.13 | 0.28 |
| *Mimosa claussenii var. claussenii* | Liau-Kang et al. 2024 | 2 | 1 | 0&4&5 | 0 | 12.90 | 19.80 | |
| *Mimosa claussenii var. claviceps* | Liau-Kang et al. 2024 | 2 | 1 | 0 | 0 | 14.30 | 23.00 | |
| *Mimosa cordistipula* | Lima et al. 2008 | 2 | | 0 | 0&6 | 21.70 | 26.10 | 1.00 |
| *Mimosa coruscocaesia* | Santos-Silva et al. 2013 | 2 | 1 | 3 | | 13.80 | 19.70 | |
| *Mimosa craspedisetosa* | Santos-Silva et al. 2013 | 2 | 1 | 3 | 0&6 | 12.20 | 15.25 | |
| *Mimosa cruenta* | Liau-Kang et al. 2024 | 2 | 1 | 5 | 2&3 | 9.54 | 14.91 | 0.16 |
| *Mimosa daleoides* | Medina-Acosta-2018 | 2 | | 4 | 6 | 31.27 | 32.54 | 1.57 |
| *Mimosa dalyi* | Santos-Silva et al. 2013 | 2 | 1 | 3 | 6 | 13.00 | 15.60 | |
| *Mimosa debilis var. debilis* | Lima et al. 2008 | 2 | 1 | 0&5 | 0 | 10.20 | 10.70 | 0.80 |
| *Mimosa dichroa* | Santos-Silva et al. 2013 | 3 | 1 | 4 | 0 | 8.90 | 10.50 | |
| *Mimosa diplotricha var. diplotricha* | Liau-Kang et al. 2024 | 2 | 1 | 3 | 2 | 25.97 | 34.06 | 1.09 |
| *Mimosa dolens var. dolens* | Liau-Kang et al. 2024 | 2 | 0 | 5 | 6 | 11.99 | 13.17 | 0.27 |
| *Mimosa dryandroides* | Liau-Kang et al. 2024 | 2 | 1 | 0&5 | 0&6 | 8.68 | 13.12 | 0.24 |
| *Mimosa elliptica* | Cruz et al. 2018 | 3 | 1 | 3 | 3 | 22.00 | 27.80 | 2.00 |
| *Mimosa ephedroides* | Caccavari 2002 | 3 | 2 | 4 | 0 | | 18.00 | |
| *Mimosa exalbescens* | Lima et al. 2008 | 3 | 1 | 0 | 0&6 | 10.50 | 13.80 | 0.50 |
| *Mimosa fiebrigii* | Santos-Silva et al. 2013 | 2 | 1 | 3 | 0&6 | 14.50 | 17.10 | |
| *Mimosa filipes* | Lima et al. 2008 | 2 | 1 | 0 | 0&6 | 20.50 | 21.70 | 1.10 |
| *Mimosa floridana* | Flores-Cruz et al. 2006 | 2 | 1 | 1 | 6 | 25.00 | 28.00 | 1.30 |

| Taxon names | Literature source | D.U. Number of grains | D.U. Outline | Pollen arrangement | Pollen ornamentation | D.U. Shorter diameter | D.U. Longer diameter | Pollen exine thickness |
|---|---|---|---|---|---|---|---|---|
| *Mimosa foliolosa var. brevibractea* | Liau-Kang et al. 2024 | 2 | 1 | 0&3 | 0 | 13.00 | 20.10 | |
| *Mimosa foliolosa var. franciscana* | Liau-Kang et al. 2024 | 2 | 1 | 0&3 | 0 | 12.50 | 20.60 | |
| *Mimosa foliolosa var. multipinna* | Liau-Kang et al. 2024 | 2 | 1 | 0&3 | 0 | 13.10 | 19.70 | |
| *Mimosa foliolosa var. pachycarpa* | Liau-Kang et al. 2024 | 2 | 1 | 3 | 0 | 13.47 | 18.02 | 0.38 |
| *Mimosa foliolosa var. paranani* | Liau-Kang et al. 2024 | 2 | 1 | 3&5 | 0&2 | 19.58 | 12.46 | 0.23 |
| *Mimosa foliolosa var. pubescens* | Liau-Kang et al. 2024 | 2 | 1 | 3&5 | 0&2 | 13.30 | 19.14 | 0.36 |
| *Mimosa foliolosa var. vernicosa* | Liau-Kang et al. 2024 | 2 | 1 | 0&4&5 | 0 | 12.12 | 18.80 | |
| *Mimosa foliolosa var. viscidula* | Liau-Kang et al. 2024 | 2 | 1 | 0 | 0 | 13.00 | 20.60 | |
| *Mimosa gemmulata* | Liau-Kang et al. 2024; Lima et al. 2008; Santos-Silva et al. 2013 | 2 | 1 | 0&4&5 | 0&6 | 15.62 | 21.14 | 1.00 |
| *Mimosa glutinosa* | Santos-Silva et al. 2013 | 2 | 1 | 3 | 0 | 12.80 | 18.50 | |
| *Mimosa guilandinae var. extensissima* | Caccavari 2002 | 5 | 1 | 1 | 0 | | 23.00 | |
| *Mimosa hapaloclada* | Santos-Silva et al. 2013 | 3 | 1 | 0 | 6 | 15.50 | 22.10 | |
| *Mimosa hebecarpa* | Santos-Silva et al. 2013 | 2 | 1 | 3 | 0 | 14.00 | 18.10 | |
| *Mimosa hexandra* | Lima et al. 2008 | 3 | | 0 | 0&6 | 10.90 | 14.60 | 0.50 |
| *Mimosa hystricina* | Flores-Cruz et al. 2006 | 2 | 1 | 0 | 6 | 24.00 | 27.00 | 1.30 |
| *Mimosa insignis* | Santos-Silva et al. 2013 | 2 | 1 | 3 | 0 | 12.50 | 16.60 | |
| *Mimosa interrupta* | Santos-Silva et al. 2013 | 2 | 1 | 3 | 0&6 | 10.50 | 17.50 | |
| *Mimosa invisa* | Buril et al. 2010; Lima et al. 2008 | 2 | 1&2 | 0&3 | 0&6 | 18.03 | 22.23 | 1.20 |
| *Mimosa invisa var. macrostachya* | Liau-Kang et al. 2024 | 2 | 1 | 3 | 0 | 16.96 | 23.79 | 0.34 |
| *Mimosa irrigua* | Lima et al. 2008 | 3 | | 0&5 | 0&6 | 13.10 | 18.50 | 1.00 |
| *Mimosa laniceps* | Liau-Kang et al. 2024 | 2 | 1 | 0 | 0 | 13.50 | 21.00 | |
| *Mimosa laticifera* | Liau-Kang et al. 2024; Barth 1973 | 3 | 1 | 3 | 6 | 15.57 | 20.40 | 0.31 |
| *Mimosa latidens* | Flores-Cruz et al. 2006 | 2 | 1 | 0 | 6 | 30.00 | 33.00 | 1.30 |
| *Mimosa lepidophora* | Lima et al. 2008 | 4 | 0&1 | 1 | 0 | 18.60 | 22.70 | 1.10 |
| *Mimosa lewisii* | Lima et al. 2008 | 2 | 1 | 0 | 0&6 | 17.10 | 23.50 | 1.10 |
| *Mimosa maguirei* | Liau-Kang et al. 2024 | 2 | 1 | 0&5 | 0 | 13.10 | 19.70 | |
| *Mimosa manidea* | Liau-Kang et al. 2024 | 2 | 1 | 0&5 | 0 | 9.98 | 15.65 | 0.15 |
| *Mimosa mensicola* | Lima et al. 2008; Santos-Silva et al. 2013 | 3 | 1 | 4&5 | 0&6 | 11.40 | 14.75 | 1.00 |
| *Mimosa misera* | Lima et al. 2008 | 2 | | 3 | 0&6 | 20.10 | 25.90 | 1.10 |
| *Mimosa modesta* | Lima et al. 2008 | 2 | | 0&5 | 0 | 9.30 | 9.90 | 0.70 |
| *Mimosa morroensis* | Lima et al. 2008 | 2 | 1 | 0&5 | 0&6 | 9.00 | 11.00 | 0.60 |
| *Mimosa myrioglandulosa* | Liau-Kang et al. 2024 | 2 | 1 | 3&4 | 0 | 11.29 | 16.75 | 0.26 |
| *Mimosa nothopteris* | Santos-Silva et al. 2013 | 2 | 1 | 1 | 0 | 13.00 | 16.50 | |
| *Mimosa obstrigosa* | Caccavari 2002 | 2 | 1 | 5 | 2 | | 10.00 | |
| *Mimosa occidentalis* | Medina-Acosta-2018 | 2 | 0&1 | 5 | 2 | 10.00 | 15.10 | 0.87 |

| Taxon names | Literature source | D.U. Number of grains | D.U. Outline | Pollen arrangement | Pollen ornamentation | D.U. Shorter diameter | D.U. Longer diameter | Pollen exine thickness |
|---|---|---|---|---|---|---|---|---|
| *Mimosa oedoclada* | Liau-Kang et al. 2024 | 2 | 1 | 3&5 | 0 | 10.88 | 16.55 | 0.54 |
| *Mimosa ophthalmocentra* | Buril et al. 2010; Lima et al. 2008; Santos-Silva et al. 2013 | 3 | 1 | 4 | 0&6 | 9.88 | 13.80 | 0.60 |
| *Mimosa paludosa* | Liau-Kang et al. 2024; Lima et al. 2008; Liau-Kang et al. 2024 | 2 | 1 | 0&5 | 0&6 | 15.50 | 22.55 | 1.00 |
| *Mimosa papposa* | Caccavari 2002 | 2 | 1 | 4&5 | 6 | | 12.00 | |
| *Mimosa paraibana* | Lima et al. 2008 | 3 | 1 | 0 | 0&6 | 10.20 | 13.80 | 0.50 |
| *Mimosa pigra* | Lima et al. 2008 | 2 | 1 | 0&3 | 6 | 18.10 | 25.70 | 1.00 |
| *Mimosa pigra var. dehiscens* | Liau-Kang et al. 2024 | 2 | 1 | 0&3 | 0&6 | 16.70 | 22.29 | 0.37 |
| *Mimosa piresii* | Liau-Kang et al. 2024 | 2 | 1 | 0&3 | 0 | 14.90 | 20.40 | |
| *Mimosa pithecolobioides* | Lima et al. 2008; Liau-Kang et al. 2024 | 4 | 1 | 1 | 0 | 16.97 | 23.17 | 0.31 |
| *Mimosa pringlei var. pringlei* | Medina-Acosta-2018 | 3 | 1 | 4 | 2 | 12.61 | 16.56 | 0.87 |
| *Mimosa prorepens* | Liau-Kang et al. 2024 | 2 | 1 | 0&5 | 0 | 11.80 | 16.90 | |
| *Mimosa pseudosepiaria* | Lima et al. 2008 | 3 | 1 | 0 | 0&6 | 10.40 | 14.90 | 0.50 |
| *Mimosa pteridifolia* | Liau-Kang et al. 2024; Santos-Silva et al. 2013 | 1&2 | 1 | 4&5 | 6 | 14.66 | 20.35 | 0.28 |
| *Mimosa pudica* | Lima et al. 2008 | 2 | 1 | 0&5 | 0 | 10.10 | 10.40 | 0.50 |
| *Mimosa quadrivalvis var. angustata* | Flores-Cruz et al. 2006 | 2 | 1 | 0 | 6 | 27.00 | 30.00 | 1.30 |
| *Mimosa quadrivalvis var. diffusa* | Flores-Cruz et al. 2006 | 2 | 1 | 0 | 6 | 28.00 | 31.00 | 1.30 |
| *Mimosa quadrivalvis var. jaliscensis* | Flores-Cruz et al. 2006 | 2 | 1 | 0 | 6 | 30.00 | 36.00 | 1.30 |
| *Mimosa quadrivalvis var. nuttallii* | Flores-Cruz et al. 2006 | 2 | 1 | 0 | 6 | 26.00 | 29.00 | 1.30 |
| *Mimosa quadrivalvis var. occidentalis* | Flores-Cruz et al. 2006 | 2 | 1 | 0 | 2 | 27.00 | 30.00 | 1.30 |
| *Mimosa quadrivalvis var. paucijuga* | Flores-Cruz et al. 2006 | 2 | 1 | 0 | 6 | 30.00 | 35.00 | 0.60 |
| *Mimosa quadrivalvis var. platycarpa* | Flores-Cruz et al. 2006 | 2 | 1 | 0 | 2 | 29.00 | 33.00 | 1.30 |
| *Mimosa quadrivalvis var. quadrivalvis* | Flores-Cruz et al. 2006 | 2 | 1 | 0 | 6 | 32.00 | 35.00 | 1.30 |
| *Mimosa quadrivalvis var. urbaniana* | Flores-Cruz et al. 2006 | 2 | 1 | 0 | 2 | 25.00 | 28.00 | |
| *Mimosa regnellii var. supersetosa* | Caccavari 2002 | 2 | | 0&3 | 6 | | 27.00 | |
| *Mimosa rhodostegia* | Liau-Kang et al. 2024 | 2 | 1 | 0&3 | 0 | 13.40 | 20.70 | |
| *Mimosa robusta* | Flores-Cruz et al. 2006 | 2 | 1 | 0 | 6 | 39.00 | 47.00 | 1.30 |
| *Mimosa rupigena* | Liau-Kang et al. 2024 | 2 | 1 | 0&3 | 0 | 13.40 | 19.50 | |
| *Mimosa scabrella* | Liau-Kang et al. 2024; Bauermann et al. 2009 | 2 | 2 | 0&5 | 8 | 9.58 | 13.49 | 0.77 |
| *Mimosa schomburgkii* | Santos-Silva et al. 2013 | 1&2 | 1 | 4&5 | 0&6 | 16.10 | 30.00 | |
| *Mimosa sensitiva* | Buril et al. 2010 | 2 | 1 | 5 | 0 | | 8.65 | |
| *Mimosa sensitiva var. sensitiva* | Lima et al. 2008 | 2 | 0 | 0&5 | 0 | 8.30 | 8.80 | 1.00 |
| *Mimosa sericantha* | Santos-Silva et al. 2013 | 2 | 1 | 3&5 | 6 | 11.90 | 23.00 | |
| *Mimosa setosissima* | Liau-Kang et al. 2024 | 2 | 1 | 0&5 | 0 | 12.10 | 17.20 | |
| *Mimosa setuligera* | Lima et al. 2008 | 2 | 1 | 5 | 0&6 | 10.90 | 12.70 | 1.00 |
| *Mimosa somnians var. somnians* | Lima et al. 2008 | 2 | 1 | 0&3 | 0&6 | 12.50 | 20.10 | 1.00 |

| Taxon names | Literature source | D.U. Number of grains | D.U. Outline | Pollen arrangement | Pollen ornamentation | D.U. Shorter diameter | D.U. Longer diameter | Pollen exine thickness |
|---|---|---|---|---|---|---|---|---|
| *Mimosa sousae* | Medina-Acosta-2018 | 2 | 0&1 | 5 | 2&3 | 10.33 | 14.67 | 1.04 |
| *Mimosa spirocarpa* | Medina-Acosta-2018 | 3 | 1 | 5 | 2&3 | 10.53 | 13.68 | 0.87 |
| *Mimosa spixiana* | Santos-Silva et al. 2013 | 2 | 1 | 0&5 | 0 | 16.00 | 18.20 | |
| *Mimosa strobiliflora* | Liau-Kang et al. 2024 | 2 | 1 | 0&3 | 6 | 13.78 | 20.16 | 0.33 |
| *Mimosa subenervis* | Lima et al. 2008 | 2 | 1 | 0 | 0&6 | 21.70 | 28.50 | 1.00 |
| *Mimosa subinermis* | Flores-Cruz et al. 2006 | 2 | 1 | 0 | 2&3 | 26.00 | 30.00 | |
| *Mimosa taimbensis* | Barth & Yoneshigue 1966 | 2 | 0 | 0 | 4 | 11.90 | 19.00 | 1.20 |
| *Mimosa tandilensis* | Caccavari 2002 | 2 | 1 | 0&5 | 6 | | 31.00 | |
| *Mimosa tenuiflora* | Buril et al. 2010; Lima et al. 2008; Santos-Silva et al. 2013; Liau-Kang et al. 2024 | 2 | | 0&5 | 0&7 | 14.06 | 20.37 | 0.51 |
| *Mimosa tetragona* | Flores-Cruz et al. 2006 | 2 | 1 | 0 | 2 | 25.00 | 30.00 | |
| *Mimosa trianae* | Santos-Silva et al. 2013 | 1&2 | 1 | | | 14.00 | 15.50 | |
| *Mimosa ulbrichiana* | Lima et al. 2008 | 2 | 0 | 0&3 | 0&6 | 20.50 | 26.80 | 1.00 |
| *Mimosa ulei var. ulei* | Liau-Kang et al. 2024 | 2 | 1 | 0&5 | 0 | 11.40 | 18.40 | |
| *Mimosa ursina* | Lima et al. 2008 | 0&1 | 2 | 2 | 0 | 16.00 | 18.15 | 1.70 |
| *Mimosa velloziana* | Lima et al. 2008 | 2 | 2 | 0&5 | 0 | 9.90 | 10.40 | 0.50 |
| *Mimosa verrucosa* | Lima et al. 2008; Santos-Silva et al. 2013 | 2 | 1 | 0 | 0&6 | 13.65 | 19.10 | 1.00 |
| *Mimosa watsonii* | Medina-Acosta-2018 | 4 | 1 | 2 | 2&3 | 14.54 | 20.70 | 0.75 |
| *Mimosa xavantinae* | Santos-Silva et al. 2013 | 2 | 1 | | | 13.00 | 14.50 | |
| *Mimosa xiquexiquensis* | Lima et al. 2008 | 2 | 2 | 0&5 | 0&6 | 13.00 | 16.60 | 0.50 |
| *Naiadendron duckeanum* | Guinet & Caccavari 1992 | 5 | 1&2 | | 4&6 | 26.30 | 36.60 | 0.90 |
| *Parapiptadenia blanchetii* | Caccavari 2002; Ribeiro et al. 2018 | 5 | 2 | | 4 | 26.30 | 31.28 | 1.00 |
| *Parapiptadenia excelsa* | Caccavari 2002 | 4 | 2 | | | | 36.00 | |
| *Parapiptadenia pterosperma* | Caccavari 2002; Cruz et al. 2018 | 5 | 0 | | 3 | 29.10 | 37.55 | 2.00 |
| *Parapiptadenia rigida* | Caccavari 2002 | 5 | 1&2 | | | | | |
| *Parapiptadenia zehntneri* | Caccavari 2002; Buril et al. 2010; Ribeiro et al. 2018 | 5 | 2 | 2 | 0&4 | 26.00 | 30.40 | 1.05 |
| *Piptadenia adiantoides* | Barduzzi et al. 2024; Caccavari 2002; Ribeiro et al. 2018 | 3&4 | 2 | 5 | 4 | 18.98 | 22.59 | 0.55 |
| *Piptadenia affinis* | Barth & Yoneshigue 1966 | 3 | 2 | 5 | 4 | 23.10 | 31.50 | 1.70 |
| *Piptadenia anolidurus* | Caccavari 2002 | 2&3 | 2 | | | | 25.00 | |
| *Piptadenia floribunda* | Caccavari 2002 | 5 | 1 | | | | 42.00 | |
| *Piptadenia fruticosa* | Caccavari 2002 | 3&4 | 2 | 2 | 0 | | 19.00 | |
| *Piptadenia gonoacantha* | Caccavari 2002 | 4 | 2 | | 3 | | 38.00 | |
| *Piptadenia irwinii* | Ribeiro et al. 2018 | 5 | 1&2 | 2 | 4 | 18.73 | 23.13 | 1.00 |
| *Piptadenia irwinii var. irwinii* | Caccavari 2002 | 4 | 2 | | 0 | | 26.00 | |
| *Piptadenia irwinii var. unijuga* | Caccavari 2002 | 4 | 2 | | 0 | | 25.00 | |

| Taxon names | Literature source | D.U. Number of grains | D.U. Outline | Pollen arrangement | Pollen ornamentation | D.U. Shorter diameter | D.U. Longer diameter | Pollen exine thickness |
|---|---|---|---|---|---|---|---|---|
| *Piptadenia killipii* | Caccavari 2002 | 3&4 | 2 | | 4 | | 20.00 | |
| *Piptadenia micracantha* | Caccavari 2002 | 4 | 2 | | 4 | | 19.00 | |
| *Piptadenia minutiflora* | Caccavari 2002 | 5 | 1&2 | | 4 | | 50.00 | |
| *Piptadenia paniculata* | Barduzzi et al. 2024 | 5 | 1&2 | | | 25.70 | 35.59 | 0.54 |
| *Piptadenia polyptera* | Caccavari 2002 | 3 | 2 | | | | 25.00 | |
| *Piptadenia retusa* | Caccavari 2002 | 4 | 2 | | | | 23.00 | |
| *Piptadenia santosii* | Caccavari 2002 | 4 | 2 | | | | 50.00 | |
| *Piptadenia stipulacea* | Caccavari 2002; Buril et al. 2010 | 4 | 1&2 | 2 | 0 | 19.30 | 23.40 | 1.00 |
| *Piptadenia trisperma* | Caccavari 2002; Cruz et al. 2018 | 3 | 2 | 1 | 3 | 21.30 | 26.25 | |
| *Pityrocarpa brenanii* | Barduzzi et al. 2024 | 4&5 | | | 4 | 27.99 | 33.76 | 0.35 |
| *Pityrocarpa inaequalis* | Caccavari 2002 | 4 | 2 | | 4&6 | | 43.00 | |
| *Pityrocarpa leptostachya* | Barduzzi et al. 2024 | 5 | 1&2 | | 6 | | | |
| *Pityrocarpa moniliformis* | Buril et al. 2010; Ribeiro et al. 2018; Guinet & Caccavari 1992 | 3 | 1&2 | 2 | 4&6 | 18.25 | 21.82 | 1.00 |
| *Pityrocarpa obliqua* | Caccavari 2002; Ribeiro et al. 2018 | 3&5 | 2 | | 0 | 26.00 | 30.60 | 1.00 |
| *Pityrocarpa obliqua subsp. brasiliensis* | Ribeiro et al. 2018 | 5 | 1&2 | | 0 | 25.50 | 32.15 | |
| *Pityrocarpa schumanniana* | Caccavari 2002 | 5 | 2 | | | | 42.00 | |
| *Senegalia nigrescens* | Kenrick & Knox 1982 | 5 | 1&2 | | 4&6 | | | |
| *Stryphnodendron adstringens* | Guinet & Caccavari 1992; Barduzzi et al. 2024 | 5 | 1&2 | | 4&6 | 23.11 | 31.68 | 0.61 |
| *Stryphnodendron cristalinae* | Guinet & Caccavari 1992 | 5 | 1&2 | | 6 | 27.00 | 34.10 | 1.05 |
| *Stryphnodendron gracile* | Guinet & Caccavari 1992 | 5 | 1&2 | | 6 | 34.10 | 30.80 | 1.00 |
| *Stryphnodendron platyspicum* | Guinet & Caccavari 1992 | 5 | 1&2 | | 6 | 21.28 | 31.85 | 0.76 |
| *Stryphnodendron polyphyllum* | Guinet & Caccavari 1992 | 5 | 1&2 | | 4&6 | 34.10 | 32.40 | 0.90 |
| *Stryphnodendron polyphyllum var. villosum* | Guinet & Caccavari 1992 | 5 | 1&2 | | 4&6 | 34.10 | 31.00 | 0.90 |
